# An ssRNA Phage Single-gene Lysis Protein Targeting MraY Reveals Convergent Evolution Across the RNA-DNA Phage Boundary

**DOI:** 10.64898/2026.11.09.765132

**Authors:** Prasanth Manohar, Karthik Chamakura, Jennifer Tran, Lorna Min, Ry Young

## Abstract

Small lytic phages of the ssRNA *Leviviricetes* and ssDNA *Microviricetes* achieve host lysis by expression of a single protein, designated as an Sgl protein. Four ssRNA Sgls have been shown to inhibit steps in peptidoglycan biosynthesis: three target MurJ, the Lipid II flippase, and one targets MurA, the committed step of the pathway. A fifth Sgl, the E protein of the canonical ssDNA microvirus ϕX174, inhibits MraY, which catalyzes the synthesis of the first lipid-linked intermediate. This inhibition requires SlyD, a cytoplasmic FKBP-type peptidyl-prolyl cis-trans isomerase. The Sgl of the *Caulobacter* ssRNA phage ϕCb5 (*Cebevirus halophobicum*) has not been characterized. Here, we present physiological evidence that Sgl^ϕCb5^ targets MraY. Moreover, we show that, despite the absence of detectable sequence similarity, Sgl^ϕCb5^ shares several features with the ϕX174 E protein, including the distribution of hydrophobic and charged residues, a proposed N-out, C-in membrane topology, and retention of lytic activity in an Sgl^ϕCb5^-LacZ fusion. Furthermore, expression of the *Caulobacter* cytoplasmic FKBP-type peptidyl-prolyl cis-trans isomerase CC2799 enhances Sgl^ϕCb5^ -mediated lysis in *E. coli*. These parallels between Sgl function in a DNA phage and an RNA phage either represent remarkably consonant convergent evolution or some evolutionary cross-over event for which there is currently no precedent.

## Introduction

Double-strand DNA and RNA phages encode multiple proteins for host lysis, including at minimum a holin with membrane hole-forming activity and an endolysin with muralytic activity. In contrast, lytic single-stranded DNA phages (Class: *Microviricetes*; called microviruses throughout) and ssRNA phages (Class: *Leviviricetes,* called leviviruses throughout) achieve host lysis by expression of a single protein, the Sgl (“single gene lysis”)(Chamakura & Young, 2019). The first three Sgls were identified in the two paradigm F-specific leviviruses, MS2 (*Emesvirus zinderi*) and Qβ (*Qubevirus durum*), and the paradigm microvirus ϕX174 (Table 1; Fig.1). In MS2, the Sgl, named L for lysis, is encoded in a 75-codon open reading frame (Orf) overlapping the end of the *coat* and beginning of the *rep* genes (Beremand & Blumenthal, 1979; Berkhout et al., 1985; Model et al., 1979). In Qβ, Sgl function is a “moonlighting” activity of the 420 residue Mat protein, designated as A_2_ for historical reasons (Karnik & Billeter, 1983; Model et al., 1979; Winter & Gold, 1983). In ϕX174, the Sgl is E, encoded in a 91-codon ORF entirely embedded in the +1-reading frame of the virion assembly gene *D* (Hutchison & Sinsheimer, 1966; Maratea et al., 1985a; Young & Young, 1982). The Sgls A_2_ and E have been studied in detail and found to be specific inhibitors of conserved enzymes of the Lipid II biosynthesis pathway (Bernhardt et al., 2000, 2001). A_2_ inhibits MurA, which catalyzes the committed step of the pathway, and E inhibits MraY, which catalyzes the formation of Lipid I, the first lipid-linked intermediate. Stable inhibitory complexes have been revealed by cryo-electron microscopy (Cui et al., 2017; Orta et al., 2023). In both cases, the expression of the Sgl first causes a blockade in peptidoglycan (PG) biosynthesis, as assessed by following the incorporation of radiolabelled diaminopimelate into PG, before lysis (Antillon et al., 2024). In both cases, the intoxicated cell forms characteristic lytic morphologies featuring large, relatively stable septal blebs that are indistinguishable from morphologies observed after treatment with cell-wall antibiotics. The septal-bleb morphology and PG biosynthesis blockade together have been established as phenotypic markers of “Type I” Sgls (Antillon et al., 2024). In addition to E and A_2_, three more Type I Sgls have been identified: LysM, from coliphage M, Sgl^PP7^ from the *Pseudomonas* phage PP7, and Sgl^CJ3^ (Kohga et al., 2025; Li et al., 2026). The latter Sgl was found in a leviviral genome, Changjiang3, derived from a metagenome study (Chamakura et al., 2020). All three target the Lipid II flippase, MurJ. L and the Sgls from the coliphage Hgal and the *Pseudomonas* phage PRR1 have been grouped into Type II, on the basis of having no effect on PG biosynthesis, as determined by mDAP labelling, and causing a lytic morphology characterized by multiple, disperse and unstable blebs (Antillon et al., 2024). L, as the founding member of Type II, has been studied by genetic analysis. The N-terminal half of L is highly basic and dispensable for lysis, whereas a central hydrophobic domain is sensitive to conservative mutation (Supp. Fig. 1). Although lacking any sequence similarity to L, the other Class II Sgls have a similar distribution of charged and hydrophobic residues but have not been subject to genetic analysis (Berkhout et al., 1987; Chamakura, et al., 2017a; Chamakura et al., 2017b).

**Figure 1:**
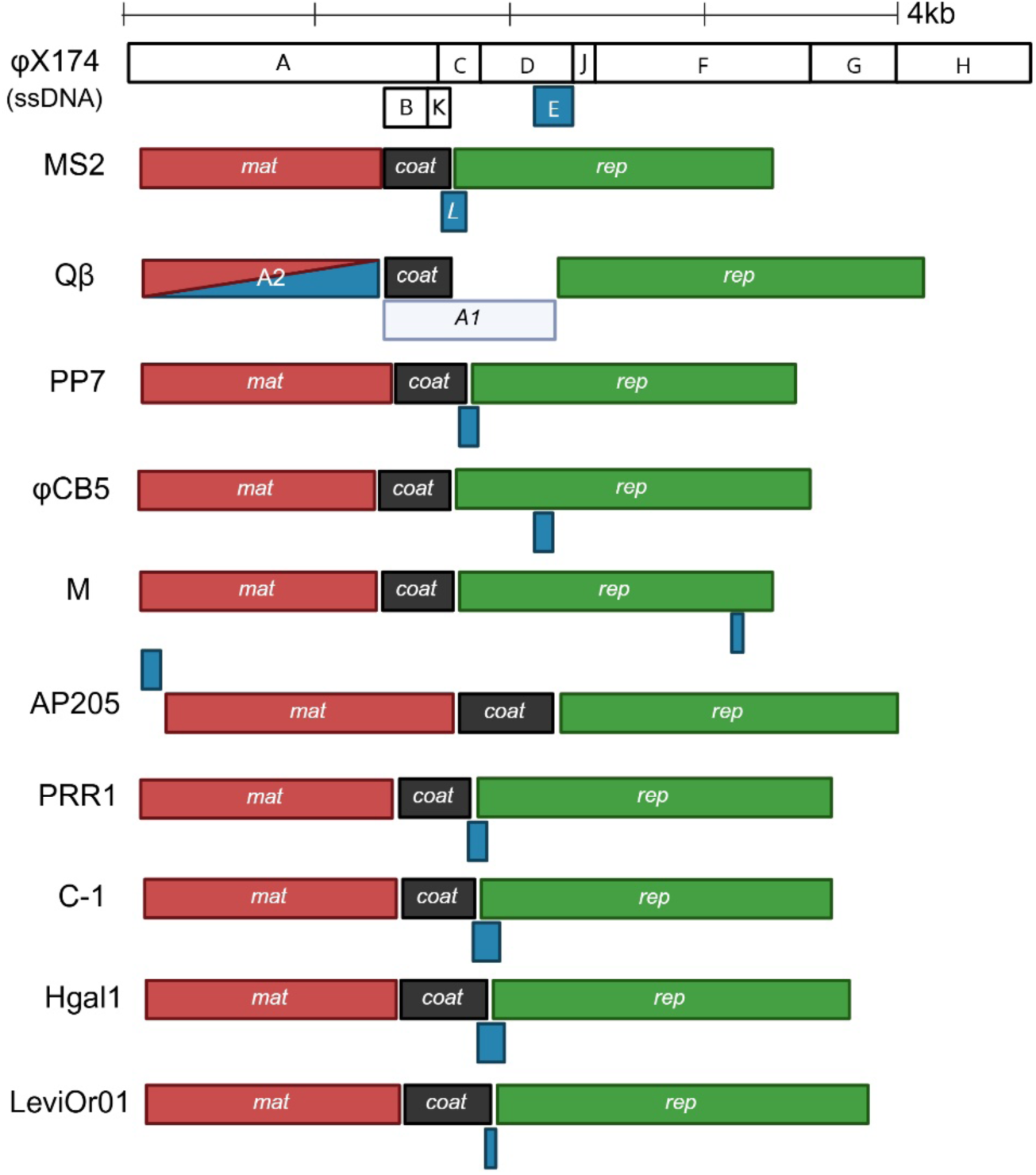
*sgl* genes in ssDNA (ϕX174) and ssRNA (MS2, Qβ, PP7, ϕCb5, M, AP205, PRR1, C-1, Hgal1, LeviOr01) phages. ssRNA phages have three core conserved genes in the same order: *mat* (red), *coat* (black), and *rep* (green). These phages also encode a single-gene lysis protein, or Sgl (blue). The Sgl is encoded in an alternative reading frame embedded in one of the three core genes or in an intergenic space. In Qβ, the Mat protein has a dual function, serving also as the Sgl.

**Table 1:** Single-Gene Lysis (Sgls) proteins of ssRNA and ssDNA phages.

| **Phage<sup>1</sup>** | **Pilus** | **Host** | **Sgl Protein** | **Type<sup>2</sup>** | **Sgl target** | **Sgl size (aa)** |
|---|---|---|---|---|---|---|
| <sup>\$</sup>MS2 | F | *E. coli* | L | II | ? | 75 |
| <sup>#</sup>Qb | F | *E. coli* | A<sub>2</sub> (Mat) | I | MurA | 420 |
| M | RIP69 (IncM) | *E. coli* | LysM | I | MurJ | 37 |
| Hgal | R27 (IncH) | *E. coli* | Sgl<sup>Hgal</sup> | II | ? | 65 |
| C-1 | RA1 (IncC) | *E. coli* | Sgl<sup>C-1</sup> | ? | ? | 65 |
| PRR1 | R1822 (IncP-1) | *P. aeruginosa* | Sgl<sup>PRR1</sup> | II | ? | 54 |
| PP7 | Polar type IV | *P. aeruginosa* | Sgl<sup>PP7</sup> | I | MurJ | 55 |
| LeviOr01 | Polar type IV | *P. aeruginosa* | Sgl<sup>Levi</sup> | ? | ? | 95 |
| AP205 | Type IV twitching | *Acinetobacter* | Sgl<sup>AP205</sup> | ? | ? | 54 |
| ϕCb5 | Polar (TAD) | *C. vibrioides* | Sgl<sup>ϕCb5</sup> | I | **MraY** | 136 |
| CJ3 | ? | *?* | Sgl<sup>CJ3</sup> | I | MurJ | 103 |
| *ϕX174 | N/A | *E. coli* | E | I | MraY | 91 |

<sup>\$</sup>Other MS2-like plaque-forming phages carrying L-like lysis proteins are fr (71 aa), M12 (75 aa), GA (63 aa), R17 (61 aa), ZR (75 aa), JP501 (75 aa), and KU1 (64 aa). <sup>#</sup>Qb-like plaque-forming phages with dual function A<sub>2</sub> protein (mat/lysis) include MX1 (421 aa), SP (576 aa), NL95 (576 aa), and M11 (588 aa). \*ϕX174 is a ssDNA phage.

The Leviviruses that have been isolated as plaque-forming phages have remarkably little diversity, despite MS2 and Qβ having been discovered decades ago. Although thousands of leviviral genomes have recently been extracted from environmental transcriptomes, fewer than 50 plaque-forming Leviviruses have been isolated and sequenced, of which most were F-specific phages of the MS2 or Qβ families (Friedman et al., 2009, 2012). Including MS2 and Qβ to represent all the F-specific phages, there are only 10 distinct ssRNA phages (Table 1 and Fig.1). As noted above, two of the remaining 8 have Type I Sgls targeting MurJ and two have Sgls experimentally determined to be Type II (Antillon et al., 2024). No *sgl* gene has been experimentally determined for the *Pseudomonas* phage LeviOr01 (Pourcel et al., 2017). Of the remaining three, the coliphage C-1 and *Acinetobacter* phage AP205 have Sgls that resemble L in terms of distribution of hydrophobic and charged residues, and one, the Sgl of the *Caulobacter* phage ϕCb5 (*Cebevirus halophobicum*), does not (Supp. Fig. 1). Here, we present studies aimed at defining whether the Sgl of ϕCb5 has Type I or Type II lytic function and, if the former, which step in the PG biosynthesis pathway it targets. The results have implications for Sgl function, regulation and evolution.

## Results

### Predicted topology and function of Sgl ϕCb5

The ϕCb5 Sgl was originally identified by Kazaks et al. (2011) by scanning the ϕCb5 genome for ORFs (open reading frames) with a predicted TMD (transmembrane domain), using the TMHMM 2.0 website (Möller et al., 2001). This revealed a 136-codon CDS (coding sequence) (nt 2284 – 2694) with a legitimate Shine-Dalgarno (AGGA, nt 2273 - 2276) upstream of a UUG start codon (Fig.1, Fig.2B; Table 1). The CDS was cloned into an unspecified pBAD vector and shown to support inducible stoppage of growth and partial reduction of culture mass (see Fig.2 of Kazaks *et al*.). The CDS, annotated then as *LP* for lysis protein, is embedded in the +1-reading frame of the *rep* gene. It was found to have two TMDs: TMD1 occupying residues 7-31 with N-in, C-out topology and TMD2 encompassing residues 89-105, with N-out, C-in topology (Supp. Fig. 1). When we re-analyzed the LP sequence with the same software, we found that while the confidence level for the TMD1 prediction was 100%, the putative TMD2 sequence registered at less than 40% and thus failed the cut-off for TMD prediction (Supp. Fig. 2). However, when we re-analyzed the LP sequence with TOPCONS, which surveys a variety of TMD prediction tools to generate a consensus (Supp. Fig. 3), which in this case was the single N-terminal TMD with N-in, C-out topology. Finally, we used DeepTMHMM, the newest TMD-prediction program, featuring deep learning methodology (Hallgren et al., 2022a). Surprisingly, DeepTMHMM predicts LP to be a soluble cytoplasmic protein with no TMDs (Supp. Fig. 2). This is unambiguously incorrect, considering the hydropathicity of the putative TMD1 sequence. We suspected that the downstream sequence designated as TMD2 in the original annotation was confounding the analysis. Indeed, DeepTMHMM analysis of an LP sequence with the downstream sequence deleted generated a high-confidence N-terminal TMD but with the opposite topology: N-out, C-in (Supp. Fig. 2). This is a clear indication that DeepTMHMM and its predecessor, TMHMM, are susceptible to dysfunction derived from the presence of downstream oligopeptide sequences of ambiguous hydropathic character (Supp. Fig. 2). In any case, it seems safe to conclude from simple inspection that LP has an N-terminal TMD, as do four other type I Sgls: E, Sgl^M^, Sgl^CJ3^ and Sgl^PP7^ (Fig. 1). Of these, the canonical Sgl E has striking parallels with LP not only in terms of the N-terminal TMD but also in terms of the distribution of charged, hydrophobic and prolyl residues, despite there being no detectable sequence similarity between the two proteins. Both have extreme N-terminal oligopeptide sequences with a single cationic and a single anionic residue. Net charge upstream of a TMD is strongly determinative of TMD topology (Maratea et al., 1985a; Young & Young, 1982). Since E has been experimentally determined to have N-out, C-in topology, it seemed likely that LP is similarly disposed. This parallelism extends to the properties of the predicted TMD sequence, both of which extend 25 residues and are leucine-rich (10 residues/25). Further, both have a C-terminal domain rich in charged residues (17 of 57 residues for E; 19 of 105 residues for LP) and especially basic (13 R/K, net charge +10 for E; 14 R/K, net charge +9 for LP). Moreover, both sequences are rich in prolines (5 of 91 for E; 7 of 136 for LP). Taking all these elements into account, we propose that LP has the same topology as E and also targets MraY as a Type I Sgl. We will refer to LP as Sgl^ϕCb5^ henceforth.

**Figure 2:**
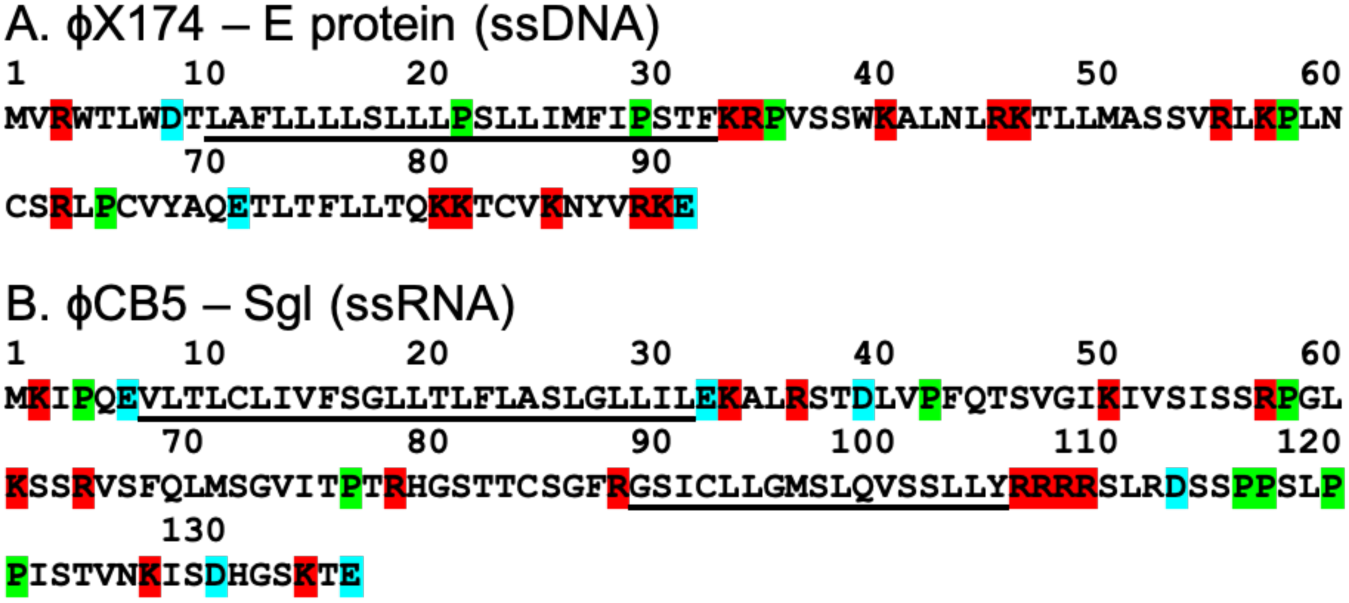
Comparison of E protein (ϕX174) (A) and Sgl^ϕCb5^ (B). The experimentally supported N-terminal transmembrane segment of Sgl^ϕCb5^ is indicated. Sgl^ϕCb5^ adopts an N-out, C-in topology. Acidic and basic residues are blue and red, respectively. Putative membrane domains are underlined.

**Figure 3:**
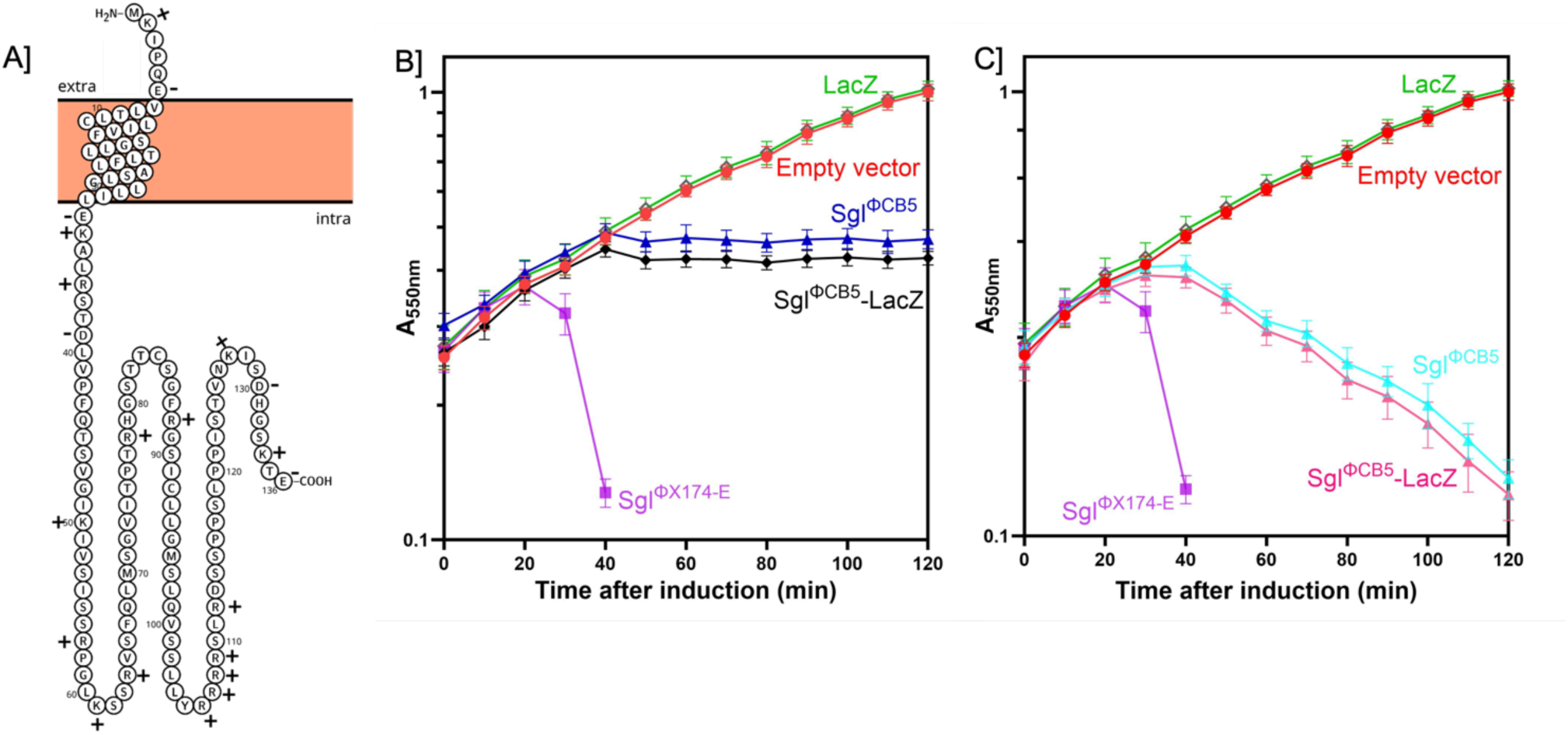
Predicted membrane topology of Sgl^ϕCb5^. (**A**) Schematic representation of the predicted membrane topology of Sgl^ϕCb5^ (136 amino acids), showing a single N-terminal transmembrane domain and a proposed N-out, C-in orientation. The short N-terminal region is positioned on the periplasmic side of the membrane, whereas the longer C-terminal region extends into the cytoplasm. (**B**) Growth curves of TB28 cultures expressing full-length *sgl^ϕCb5^*and the *lacZ* fusion. Empty vector (red), pBAD24-*lacZ* control (green), pBAD24 *sgl^ϕX174-E^-lacZ* control (purple), pBAD24 *sgl^ϕCb5^* (blue), pBAD24 *sgl^ϕCb5-^lacZ* (black). (**C**) Growth curves of TB28 Ω *mraY_Cc_*-kan cultures expressing Sgl^ϕCb5^ and the lacZ fusion. Empty vector (red), pBAD24-*lacZ* control (green), pBAD24 *sgl^ϕX174-E^-lacZ* control (purple), pBAD24 *sgl^ϕCb5^* (sky blue), pBAD24 *sgl^ϕCb5-^lacZ* (pink). The data represent the mean ± standard deviation of three independent experiments.

### Lytic activity in *E. coli*

Initially, we aimed to predict the topology of Sgl^ϕCb5^. The original studies on E topology revealed that the entire cytoplasmic domain could be replaced by a full-length β-galactosidase domain (Maratea et al., 1985b). Remarkably, the deletion-substitution *E-lacZ* retained both lytic function and full β-galactosidase activity. We constructed the full-length Sgl^ϕCb5^-LacZ using a Gly-Ser linker, and found that the Sgl^ϕCb5^-LacZ fusion protein retained the same partial lytic function in the host with the wt MraY. Moreover, it was fully lytic in an isogenic host in which the *E. coli mraY* gene was replaced by MraY_cc_ (Fig.3B). This is yet another parallel between E and Sgl^ϕCb5^ and supports the N-out, C-in topology, based on the actual fusion design and localization evidence (Fig.3A).

To characterize the Sgl^ϕCb5^, we cloned its gene into an arabinose-inducible vector plasmid. In a bulk culture of an *E. coli* MG1655 transformant, we observed growth arrest at ∼30 min after induction, indicating partial lytic function (Fig.4A). Phase-contrast microscopy revealed cells with single blebs, mostly at the septum or at a single pole (Fig.5). We considered this to be suggestive of Type I Sgl function, possibly reflective of a weak interaction between the Sgl^ϕCb5^ and the *E. coli* MraY. To test this notion, similar inductions were performed with plasmids supplying various MraY proteins in trans to the Sgl. Fig.4B shows that supplying MraY_Ec_ (*E. coli*) and MraY_Cc_ (*C. crescentus*) also resulted in growth inhibition, albeit with a slight delay that probably reflects the higher copy number of the target MraY. In contrast, supplying the MraY_Bs_ from *B. subtilis* abolishes the growth inhibition. MraY_Bs_ are significantly more diverged from the MraYs of *E. coli* and *Caulobacter* (Supp. Fig. 4) and thus presumably complement the defect in Lipid I supply caused by the inhibition of MraY_Ec_ and MraY_Cc_. This is strong evidence that Sgl^ϕCb5^ targets MraY. Moreover, the growth inhibition phenotype of Sgl^ϕCb5^ was also improved into an unambiguous lytic phenotype when we used as a host an isogenic construct in which the *E. coli mraY* gene was deleted and the *mraY* gene of *Caulobacter* supplied on a low-copy plasmid (Table 2; Fig.4C). In addition, phase contrast microscopy revealed both septal and polar single-bleb morphologies (Fig.5C, C1), further supporting the notion that Sgl^ϕCb5^ belongs to the Type I group of Sgls founded by E itself.

**Figure 4:**
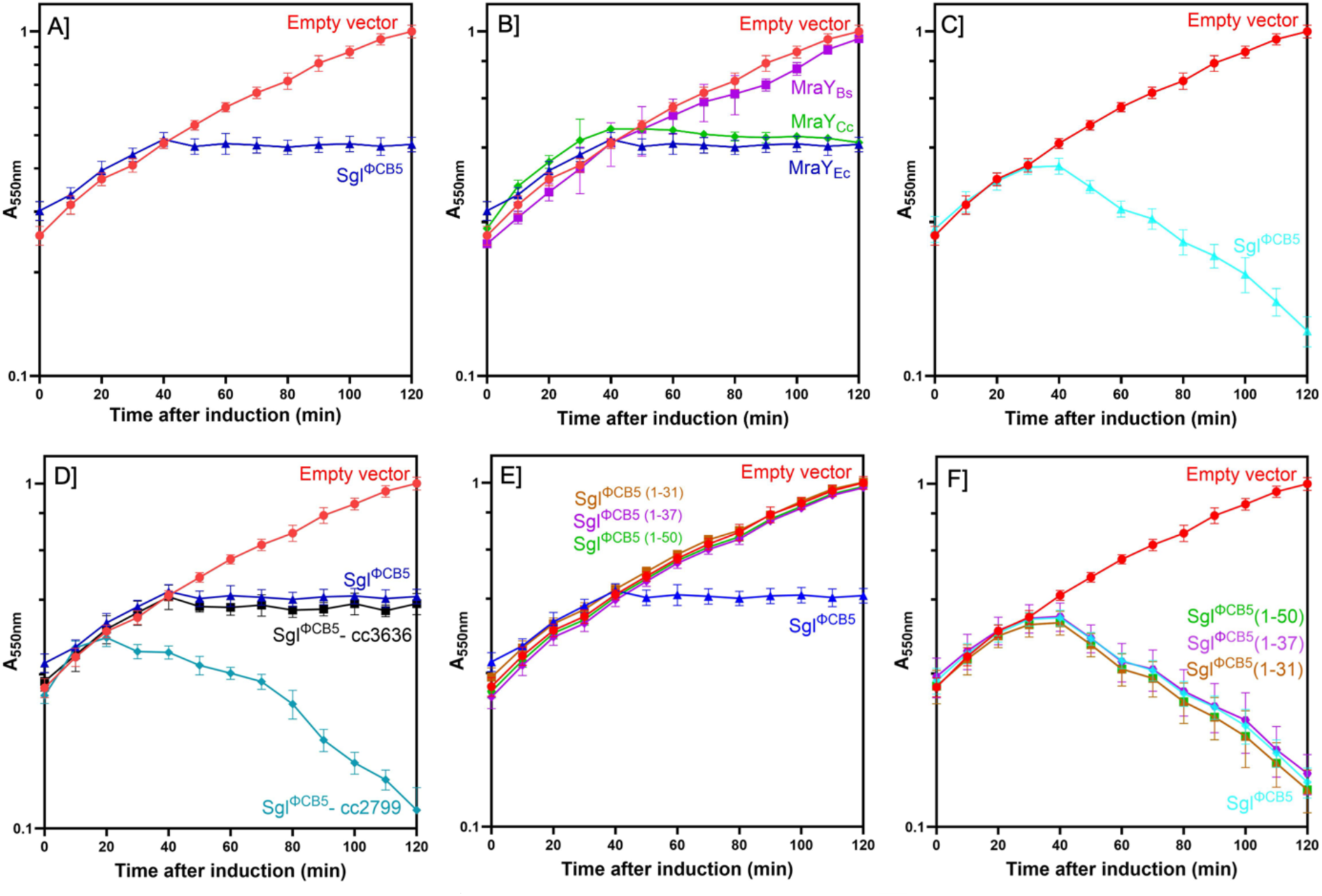
Sgl^ϕCb5^ -mediated lysis in *E. coli* MG1655. Sgl^ϕCb5^ was cloned into pBAD24 and induced in TB28 and TB28 Ω *mraY_Cc_*-kan backgrounds at 37°C. Cultures were grown to A550 = ∼0.2 and induced with arabinose at a final concentration of 0.4% (wt/vol). (**A**) Lysis profile of Sgl^ϕCb5^ in the wild-type TB28. Empty vector (red) and pBAD24 *sgl^ϕCb5^*(blue). (**B**) Lysis profile of Sgl^ϕCb5^ in the wild-type TB28 co-transformed with pBAD24 *sgl^ϕCb5^* and a compatible plasmid (pZA32) expressing MraY orthologs (MraY_Cc_ – *C. crescentus* and MraY_Bs_ – *B. subtilis*). Empty vector (red), pBAD24 *sgl^ϕCb5^* + pZA32 *mraY_Ec_* (blue), pBAD24 *sgl^ϕCb5^* + pZA32 *mraY_Cc_* (green) and pBAD24 *sgl^ϕCb5^* + pZA32 *mraY_Bs_* (purple). (**C**) Lysis profile of Sgl^ϕCb5^ in TB28 Ω *mraY_Cc_*-kan. Empty vector (red) and pBAD24 *sgl^ϕCb5^* (blue). (**D**) Lysis profile of Sgl^ϕCb5^ in the wild-type TB28 co-transformed with pBAD24 *sgl^ϕCb5^* and compatible plasmid (pZA32) expressing *C. crescentus* FKBP-type PPlases CC2799 and CC3636. Empty vector (red), pBAD24 *sgl^ϕCb5^* + pZA32 empty (blue), pBAD24 *sgl^ϕCb5^* + pZA32 CC2799 (sky blue) and pBAD24 *sgl^ϕCb5^* + pZA32 CC3636 (black). (**E**) Lysis profile of truncated Sgl^ϕCb5^ in the wild-type TB28. Empty vector (red), pBAD24 *sgl^ϕCb5^* (blue), pBAD24 *sgl^ϕCb5^* ^(1-31)^ (brown), pBAD24 *sgl^ϕCb5^* ^(1-37)^ (purple) and pBAD24 *sgl^ϕCb5^* ^(1-50)^ (green). (**F**) Lysis profile of truncated Sgl^ϕCb5^ in TB28 Ω *mraY_Cc_*-kan. Empty vector (red), pBAD24 *sgl^ϕCb5^* (blue), pBAD24 *sgl^ϕCb5^* ^(1-31)^ (brown), pBAD24 *sgl^ϕCb5^* ^(1-37)^ (purple) and pBAD24 *sgl^ϕCb5^* ^(1-50)^ (green). The data represent the mean ± standard deviation of three independent experiments.

**Figure 5:**
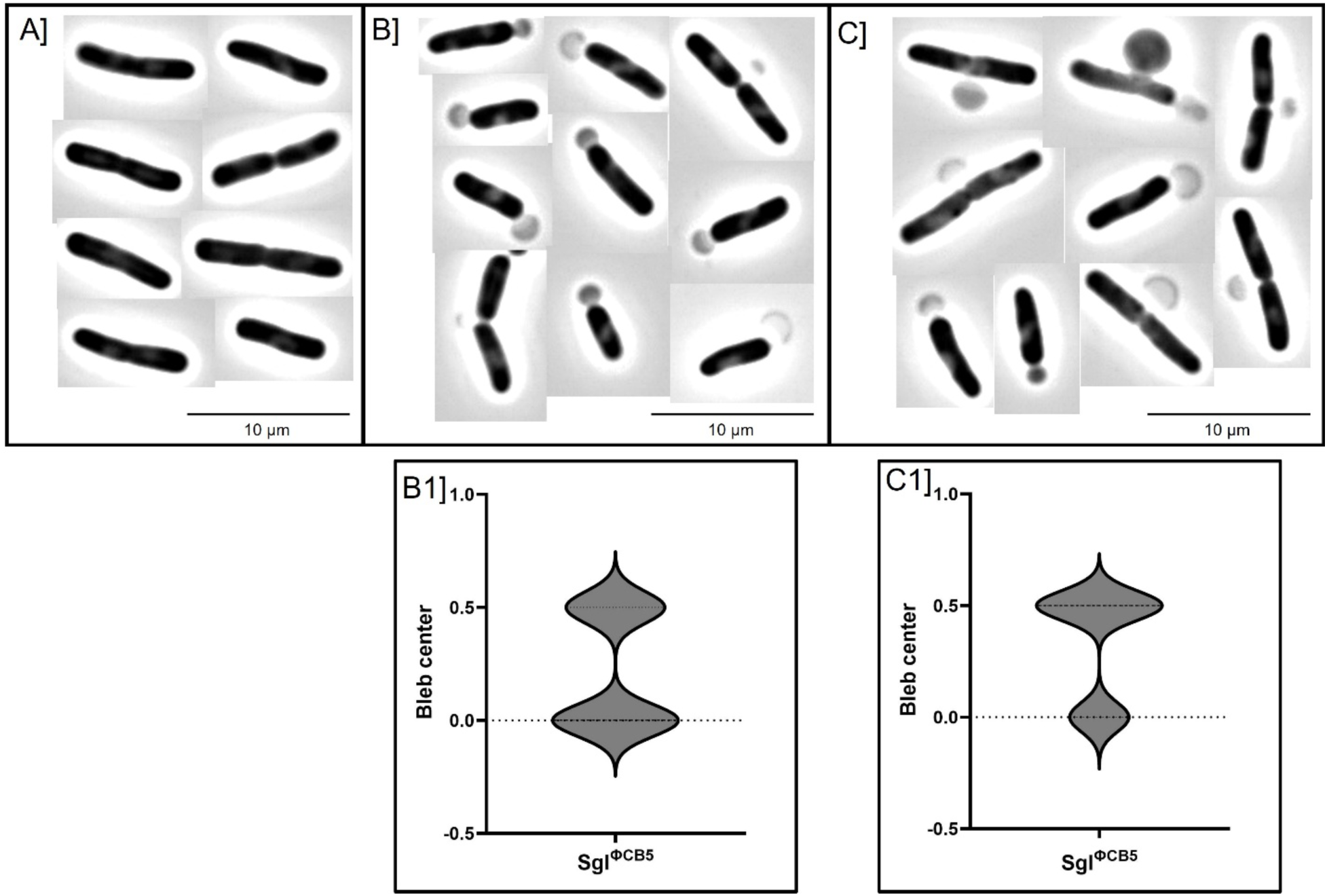
Sgl^ϕCb5^ portrays type I lysis morphology. (**A**) Phase-contrast images of *E. coli* MG1655 (TB28) control, (**B**) Bleb formation after the induction of Sgl^ϕCb5^ in *E. coli* TB28 with native MraY (MraY_Ec_), (**C**) Bleb formation after the induction of Sgl^ϕCb5^ in *E. coli* TB28 Ω *mraY_Cc_* with *C. crescentus* MraY (MraY_Cc_). Analysis of lesions caused by Sgl^ϕCb5^ in TB28 (**B1**) and TB28 Ω *mraY_Cc_* (**C1**). The y-axis represents the location of the bleb along the cell length, with 0.0 representing the poles of the cell and 0.5 representing the middle of the cell. Measurements of cells (n=25) were done using the measurement tool in ImageJ. The violin plots were created using GraphPad Prism 10.2.2. To minimize bias in the selection of cells with blebs for the analysis, all cells showing blebs in an image were measured.

**Table 2:** Bacteriophage, bacterial strains and plasmids used in this study.

| Name | Genotype and features | Source |
|---|---|---|
| **Bacteriophage** |  |  |
| ϕCb5 | Wild type ssRNA phage ϕCb5 | Laboratory stock (from K. Tars) |
| **Bacterial strains** |  |  |
| MG1655 | *E. coli* K-12 *F<sup>-</sup> ilvG rfb50 rph1* | Laboratory stock |
| TB28 (RY34010) | MG1655 *ΔlacIZYA* | (Bernhardt & De Boer, 2003) |
| RY3316 | MG1655 pKOMY3ΔmraY | (Zheng et al., 2008) |
| RY18059 | RY3316 *recA* | Laboratory stock |
| RY44251 | RY18059-derivative Ω MraY<sub>Cc</sub>-kan | (Zheng et al., 2008) |
| **Plasmids** |  |  |
| pBAD24 | pBR322-derived plasmid with P<sub>BAD</sub> promoter; *araC bla*. Amp<sup>R</sup> | Addgene ID:1845 |
| pBAD24_Sgl<sup>ϕCb5</sup> | pBAD24 vector expressing Sgl<sup>ϕCb5</sup> with ATG start codon; AGGAGG S-D sequence | This study |
| pBAD24_Sgl<sup>ϕCb5</sup>_LacZ | pBAD24 vector expressing Sgl<sup>ϕCb5</sup> fused (in-frame) with full-length LacZ | This study |
| pBAD24_Sgl<sup>ϕX174</sup>-E 35 _LacZ | pBAD24 vector expressing truncated (1-35) Sgl<sup>ϕX174</sup> fused (in-frame) with full-length LacZ | This study |
| pBAD24_Sgl<sup>ϕCb5</sup> (1-31) | pBAD24 vector expressing truncated Sgl<sup>ϕCb5</sup> *Δ(32-136)* | This study |
| pBAD24_Sgl<sup>ϕCb5</sup> (1-37) | pBAD24 vector expressing truncated Sgl<sup>ϕCb5</sup> *Δ(38-136)* | This study |
| pBAD24_Sgl<sup>ϕCb5</sup> (1-50) | pBAD24 vector expressing truncated Sgl<sup>ϕCb5</sup> *Δ(51-136)* | This study |
| *pZA32 | p15A ori, pLtetO-1 promoter, Cm<sup>R</sup> | Laboratory stock |
| pZA32_cc2799 | pZA32 expressing *C. crescentus* CC2799; Cm<sup>R</sup> | This study |
| pZA32_cc3636 | pZA32 expressing *C. crescentus* CC3636; Cm<sup>R</sup> | This study |
| pZA32_*mraY<sub>Ec</sub>* | pZA32 expressing *E. coli* MraY; Cm<sup>R</sup> | This study |
| pZA32_*mraY<sub>Bs</sub>* | pZA32 expressing *B. subtilis* MraY. Cm<sup>R</sup> | This study |
| pZA32_*mraY<sub>Cc</sub>* | pZA32 expressing *C. crescentus* MraY; Cm<sup>R</sup> | This study |
| pZA22-Kan | pZA32 derivative by replacing the *cat* gene (Cm<sup>R</sup>) with a *Kan<sup>R</sup>* cassette | Laboratory stock |
| pZA22-Kan_*mraY<sub>Cc</sub>* | pZA22 expressing *C. crescentus* MraY; Kan<sup>R</sup> | This study |

\*\*The pZA32 empty vector map was provided in Supp. Fig. 12.

### A cytoplasmic PPIase enhances Sgl^ϕCb5^-mediated lysis

E-mediated lysis requires the function of SlyD, a cytoplasmic PPIase; in its absence, E is rapidly degraded (Bernhardt et al., 2002). This likely reflects the proline-rich, highly cationic C-terminal domain of E, which is predicted to be unstructured (Orta et al., 2023). Indeed, *slyD*-dependence is abolished if the C-terminal proline-rich domain is deleted or replaced by a heterologous folding domain (Bernhardt et al., 2002b; Roof et al., 1994; Zheng et al., 2008). We hypothesized that the partial lytic function of Sgl^ϕCb5^ in *E. coli* could be improved by a similar PPIase from *Caulobacter*. PPIases fall into three structurally and enzymatically dissimilar groups: the parvulins, the cyclophilins and the FKBP-like types (Anchal et al., 2021; Ünal & Steinert, 2014). Bacterial cells have at least one of each type in each compartment. Among 9 total PPIases in *E. coli*, SlyD is one of three FKBP-like PPIases in the *E. coli* cytoplasm. In *Caulobacter* (*C. vibrioides* CB15, NC_002696.2), there are 9 PPIases but only two FKBP-types, CC3636 ([AAK25598.1](https://www.ncbi.nlm.nih.gov/protein/13425390)) and CC2799 ([AAK24763.1](https://www.ncbi.nlm.nih.gov/protein/13424401)) (Supp. Fig. 5). The former has a signal sequence, predicted with 99% confidence by SignalP, indicating that CC2799 is the only FKBP-like PPIase in the cytoplasm. Thus, we hypothesized that CC2799 would enhance the lytic activity of Sgl^ϕCb5^ if the two proteins were expressed together in *E. coli*. As expected, expression of CC2799 enhanced the lytic function of Sgl^ϕCb5^ in *E. coli* (Fig.4D). Expression of CC3636 had no effect, which is not surprising since it is likely exported to the periplasm. We tested truncation alleles of *sgl^ϕCb5^*to see if the PPIase dependency would no longer apply, as observed for E; however, *sgl* ϕCb5 C-terminal deletion alleles were completely non-functional in *E. coli* (Fig.4E). Interestingly, the same deletion alleles were fully lytic in the *mraY_Cc_* substitution host (Fig.4F). The simplest interpretation is that the Sgl-MraY interaction is much stronger with MraY_Cc_ than with *E. coli* MraY. We also confirmed that the C-terminal domain of Sgl^ϕCb5^ is necessary for its interaction with CC2799 in the *mraY_Cc_* substitution host (Supp. Fig. 6).

### Bioinformatic analysis

AlphaFold 3.0 (AF3) prediction of Sgl^ϕCb5^ revealed a predominantly low-confidence structure, with extensive regions predicted to be disordered. To investigate potential interactions with its proposed host target, Sgl^ϕCb5^ was modeled with MraY from *E. coli* and *C. crescentus*. Although the resulting models suggested possible spatial associations between Sgl^ϕCb5^ and MraY, the predicted protein-protein interfaces exhibited low confidence, preventing reliable identification of specific interaction sites (Supp. Fig. 7). We further examined whether the FKBP-like protein CC2799 could be accommodated in a predicted complex with Sgl^ϕCb5^ and MraY. The ternary models containing *C. crescentus* MraY exhibited higher interface predicted template modeling (ipTM) scores (0.34-0.38) than those containing *E. coli* MraY (0.12-0.17). Nevertheless, the overall interface confidence remained low, and the models did not support a well-defined arrangement of the three proteins (Supp. Fig. 8). The predicted binary complex of Sgl^ϕCb5^ and the FKBP-like CC2799 positioned the C-terminal region of Sgl^ϕCb5^ near the folded domain of CC2799 (Supp. Fig. 9). However, the predictions exhibited low interface confidence (ipTM scores of 0.16-0.23). Thus, the AF3 models were considered exploratory structural predictions and were not used to assign residue-level interactions or infer direct protein binding.

## Discussion

### The dominance of Type I lysis

We have provided compelling physiological evidence that Sgl^ϕCb5^ targets MraY. As shown in Table 1, there are 12 Sgls, not including the Sgls from the 50 F-specific phages that are close relatives of MS2 and Qβ. As of this work, Sgl^ϕCb5^ becomes the 6^th^ Type I Sgl overall. Thus, half of the functionally identified Sgls achieve lysis by inhibiting a specific step in the Lipid II pathway segment of peptidoglycan biosynthesis. It is puzzling that, to date, no Sgl has been found to target PG biosynthesis steps beyond the MurJ flippase step, considering the existence of the antibiotic-sensitive PG biosynthesis enzymes in the periplasm. It is also worth noting that there are now five examples of monotopic membrane protein Sgls with Type I function. In all five cases, the single TMD acts by occluding the hydrophobic binding pocket for the undecaprenol moiety of the substrate (see below). In one case, the canonical Sgl E, biochemical analysis found that the inhibition of MraY is affected in a non-competitive manner (Zheng et al., 2009). This functional convergence probably reflects the constraints on evolving an inhibitor protein which has both a specific interaction surface and the ability to achieve a three-dimensional fold. Because of the lack of coding space in the Leviviruses and Microviruses, all five are encoded by reading frames embedded out of frame inside an essential gene. It is presumably relatively easy to evolve a transmembrane domain, where all that is required is ∼16 residues of sufficient hydrophobic character to achieve insertion in the bilayer as an alpha helix. Moreover, three have N-in topology and target MurJ and two have N-out topology and target MraY. It seems prudent to explore whether these trends could facilitate functional analysis of other as-yet unclassified Sgls discovered in our screening of candidates identified by bioinformatic analysis of “dark matter” ssRNA genomes (Chamakura et al., 2020).

### Convergent evolution across the ssRNA and ssDNA phage worlds

Despite the absence of detectable sequence similarity between the canonical ϕX174 E lysis protein and Sgl^ϕCb5^, the two proteins exhibit striking parallels in the distribution of hydrophobic, anionic, cationic, and proline residues. These similarities extend to their N-terminal TMDs and N-out, C-in membrane topology (Fig. 2B). Both proteins are also associated with MraY-dependent lysis. In the ϕX174 E lysis pathway, the FKBP-like peptidyl-prolyl cis-trans isomerase (PPIase) SlyD is required for efficient lysis. Structural analysis has revealed a heterohexameric inhibitory complex, designated YES, comprising two copies each of MraY, E, and SlyD (Orta et al., 2023). Within this complex, the N-terminal TMD of E occupies a cleft between TMDs 5 and 9 of MraY, obstructing access of the lipid substrate, while an adjacent cytoplasmic helix interferes with active-site closure. Given the similarities between E and Sgl^ϕCb5^ in terms of the distribution of charged, hydrophobic and proline residues, we hypothesize that the N-terminal TMD of Sgl^ϕCb5^ may interact with the same substrate-binding cleft of MraY. However, AlphaFold 3 predictions did not confidently resolve the Sgl^ϕCb5^-MraY interface, and the details of the interface remain to be determined. Nevertheless, the final and most striking parallel derives from the ability of the cytoplasmic FKBP-like PPIase CC2799 of *Caulobacter* to promote the lytic function of Sgl^ϕCb5^, in a manner depending on the presence of the otherwise dispensable C-terminal domain of the Sgl. These results suggest that YES complex (E/MraY/SlyD) formed in the ssDNA phage ϕX174 infections of E. coli is likely to have a parallel (Sgl^ϕCb5^/MraY_Cc_/CC2799) in the ssRNA phage ϕCB5 infections of *C. crescentus*. The simplest notion is that convergent evolution has occurred in the ssRNA and ssDNA worlds, resulting in very similar molecular detail. This seems surprising, especially considering the constraints placed on the two *sgl* genes, both reading frames being embedded in the cistron of essential phage genes (*D* in the case of ϕX174 *E* and *rep* in the case of ϕCB5). Alternatively, some unprecedented cross-over event between these two worlds may have occurred. In any case, the ability to reproduce the MraY inhibition system by porting MraYCc, Sgl^ϕCb5^ and the *Caulobacter* cytoplasmic FKBP-type PPIase into *E. coli* should allow a genetic approach to refining our understanding of the Sgl-MraY interface, an approach that proved very successful with the E/MraY_Ec_/SlyD system (Bernhardt et al., 2002; Zheng et al., 2009; Orta et al., 2023).

Finally, we were recently able to identify two Sgl^ϕCb5^ homologs (GenBank accessions MN033099 and MN034746; Supp. Fig. 10) that were found in metatranscriptomic whole-genome records. These Sgls are 71% (78% similarity) and 48% (58% similarity) identical to Sgl^ϕCb5^ (Supp. Fig. 11). This is the first time a Type I Sgl identified in a phage genome has been found to have homologs in metatranscriptomes, indicating that the diversity of Sgls is not boundless. In any case, the ϕCb5 family of Sgls, with three total members, is the third largest family, after the MS2 L and Qβ A_2_ Sgl families.

## Methods

### Bacterial Strains and Plasmid Construction

Bacterial strains and plasmids used in this study are listed in Table 2. Routine culturing was performed in lysogeny broth (LB) supplemented with appropriate antibiotics. The host strains used were *E. coli* DH5α and TB28. For complementation studies, strain RY18059, a derivative of RY3316 *recA* carrying a chromosomal deletion of *mraY* complemented by the temperature-sensitive plasmid pKOMY3, was used (Zheng et al., 2008). To replace the native *E. coli mraY* with *Caulobacter crescentus mraY* (*mraY_Cc_*), plasmid pZA22-MraY_Cc_ (pZA22 is a low-copy expression vector under the control of the pLtetO-1 promoter, Kan^R^) was introduced into RY18059, and cells were grown at 42°C to eliminate pKOMY3. The resulting strain, RY44251 (TB28 Ω *mraY_Cc_*-kan), was used for complementation assays.

All plasmid constructs were generated by Gibson Assembly using 20-bp homologous overlaps. The putative lysis gene (*sgl*; nt 2284–2694) from phage ϕCb5 (NC_019453.1) was cloned into pBAD24. The *mraY* genes from *E. coli* (NC_000913.3), *C. crescentus* (ABZ72797.1), and *B. subtilis* (NP_389402.2) were cloned into pZA32, a low-copy expression vector under the control of the pLtetO-1 promoter, which carries a chloramphenicol resistance (Cm^R^) (Supp. Fig. 12). Similarly, *C. crescentus* genes, *cc2799* (AAK24763.1) and *cc3636* (AAK25598.1) were cloned into pZA32.

For topology analysis, C-terminal Sgl–LacZ in-frame fusion proteins were constructed by fusing full-length Sgl^ϕCb5^ (136 aa) to full-length β-galactosidase (LacZ; 1,024 aa) through a flexible Gly-Ser (GS) linker in pBAD24. As a positive control, the N-terminal transmembrane domain of Sgl^ϕX174^-E (35 aa) was similarly fused to LacZ.

### Physiological Procedures

*E. coli* cultures carrying pBAD24-derived plasmids were grown in LB medium supplemented with 100 μg/mL ampicillin. Overnight cultures were prepared in LB containing the appropriate antibiotics (100 μg/mL ampicillin, 50 μg/mL kanamycin, or 25 μg/mL chloramphenicol).

For induction experiments, overnight cultures were diluted 1:200 into fresh medium and grown with aeration at 37°C until an optical density (A_550_) of approximately 0.2 was reached. Expression was induced with 0.4% (w/v) L-arabinose. Cell growth and lysis kinetics were monitored by measuring A_550_ at 10-min intervals. The lysis curve was plotted using GraphPad Prism 10.2.2. All the data are presented as mean ± standard deviation (SD) of at least three independent experiments with error bars.

### Microscopy

To monitor cellular morphology during lysis, samples were collected 10 min before induction and at the onset of lysis. Aliquots (3 μL) were mounted on glass microscope slides (25 × 75 mm) and covered with 22 × 22 mm coverslips. Cells were visualized using a Zeiss inverted microscope equipped with a 100× oil-immersion objective.

Images were acquired with a 100-ms exposure time, and imaging sessions were limited to 20 min to minimize phototoxicity and imaging artifacts. Image analysis was performed using ImageJ (Schneider et al., 2012). Bleb diameters were measured in triplicate for each cell, with measurements taken from blebs located either at the cell poles or along the midcell region. GraphPad Prism 10.2.2 was used to create the violin plots.

### Bioinformatic Analyses

Genome annotations and sequence similarity searches were performed using BLASTP and PSI-BLAST available through the NCBI platform. Genome visualization and annotation editing were conducted using Artemis (Carver et al., 2012). Predicted transmembrane domains and membrane topology were analyzed using TMHMM 2.0 (Möller et al., 2001), DeepTMHMM (Hallgren et al., 2022b), and TOPCONS (Tsirigos et al., 2015). Multiple sequence alignments were generated using CLC Main Workbench and compared with homologous proteins identified through database searches.

### AlphaFold3 Structural Predictions

Structural predictions were generated using the AlphaFold 3 server (Abramson et al., 2024). Amino acid sequences were obtained from NCBI databases using the following accession numbers: ϕCb5 Sgl (NC_019453.1), MraY_Ec_ (NC_000913.3), MraY_Cc_ (ABZ72797.1), and MraY_Bs_ (NP_389402.2). We modelled binary complexes involving MraY_Ec_-Sgl^ϕCb5^, MraY_Cc_-Sgl^ϕCb5^ and Sgl^ϕCb5^-CC2799 at 1:1 stoichiometry, with one copy of each protein. Ternary complexes comprising MraY_Ec_-Sgl^ϕCb5^-CC2799 and MraY_Cc_-Sgl^ϕCb5^-CC2799 were modelled at 1:1:1 stoichiometry, with one copy of each protein. Five predicted models were evaluated for each complex using pTM, ipTM, pairwise ipTM, and predicted alignment error (PAE). For structural visualization and analysis, the predicted protein structures and complexes generated by AF3 were visualized using PyMOL (Schrödinger, LLC). Molecular representations, including cartoon and molecular surface views, were prepared to examine the spatial arrangement of Sgl^ϕCb5^ relative to MraY and CC2799. Putative interaction regions were examined by visual inspection of the predicted model. The predictions were considered exploratory and were not used to establish direct protein-protein interactions or assign experimentally validated contact residues.

## Acknowledgment

The authors would like to thank the funding supported by the National Institute of General Medical Sciences, grant no. R35GM136396 to R.Y. and by the Center for Phage Technology, which is jointly supported by Texas A&M AgriLife Research.

## Author contribution

KC conducted the original experiments identifying the host target. PM analyzed the lytic phenotypes and wrote the original draft MS. RY edited and expanded the MS and provided funding and laboratory space. JT and LM conducted the original *Caulobacter* PPIase dependency and ϕCb5 truncation experiments.

## Supplementary information

Supplementary Figure 1: Sequence alignment of L-like Single-Gene Lysis (Sgl) proteins.

Supplementary Figure 2: Problematic TMD topology prediction for Sgl^ϕCb5^.

Supplementary Figure 3: The graphical output of TOPCONS showing Sgl^ϕCb5^ topology.

Supplemental Figure 4: Alignment of MraY from *E. coli*, *C. crescentus* and *B. subtilis*.

Supplementary Figure 5: Amino acid sequences of *C. vibrioides* FKBP-type PPIase; (A) CC2799 and (B) CC3636.

Supplementary Figure 6: Effect of CC2799 expression on lysis mediated by full-length and truncated Sgl^ϕCb5^.

Supplementary Figure 7: AlphaFold 3 predictions of MraY-Sgl^ϕCb5^ complexes.

Supplementary Figure 8: AlphaFold 3 predictions of MraY-Sgl^ϕCb5^ -CC2799 ternary complexes.

Supplementary Figure 9: AlphaFold 3-predicted structural model of Sgl^ϕCb5^ in association with the FKBP-like CC2799.

Supplementary Figure 10: Comparative genome organization and distribution of charged residues in putative Sgl proteins of ϕCb5-related ssRNA phages.

Supplementary Figure 11: Multiple sequence alignment of Sgl^ϕCb5^ and two putative homologs.

Supplementary Figure 12: Genetic map of the pZA32 empty vector (2,108 bp).

