## Supplementary file for "An ssRNA Phage Single-gene Lysis Protein Targeting MraY Reveals Convergent Evolution Across the RNA-DNA Phage Boundary"

### Supplementary Figures:

```
1        10          20          30          40          50          60          70
METRFPQQSQQTPASTNRRRPFKHEDYPCRRQQRSSTLYVLIFLAIFLSKFTNQLLLSLLEAVIRTVTTLQQLLT          (MS2)
    MQQPSQPTRESTKKPVPFQHEEYPCQNQQRSSTLYVLICLAIFLSKFTNQLLASLLDLLIRIVTTLQQLLT          (fr)
                MGQKAFKHKENLCSDSQRSKRLYVWIALAIVLSDFTSIFSHWIWGLLILILRTLMDLPTFVMNV     (KU1)
            MAKPSNRRRSLREENPCRAERVPKRTVVFAVSAICLSKYINQYLANLLDLLIQCLQSLIRLLT          (AVE017)*
    MANPRRETLSVHRFIVTKSQAQFSIRIIRCIRNLLAIAGMLFLLSPIYLDKFIQIYLNAYVVPM                 (HGAL)
 MPSLTLRVSTNLPDITVRDAIHKGYLWLRKQVAVATVWLLILTFIGLSSGKLRLWFDLSVTNLSG                   (C1)
                               MKKRTKALLPYAVFIILSFQLTLLTALFMYYHYTF                   (AP205)
 MCKYSTKVDSKLTESVGQLTIRSYLWLRNILALAGLLFVILLATNHLSIAIYSP                              (PRR1)
```

**Supplementary Figure 1: Sequence alignment of L-like Single-Gene Lysis (Sgl) proteins.** MS2, fr, KU1, Hgal, C-1, AP205 and PRR1 are plaque-forming ssRNA phages, and AVE017 is a metatranscriptomic sequence. Acidic and basic residues are blue and red, respectively. Underline denotes TMD.

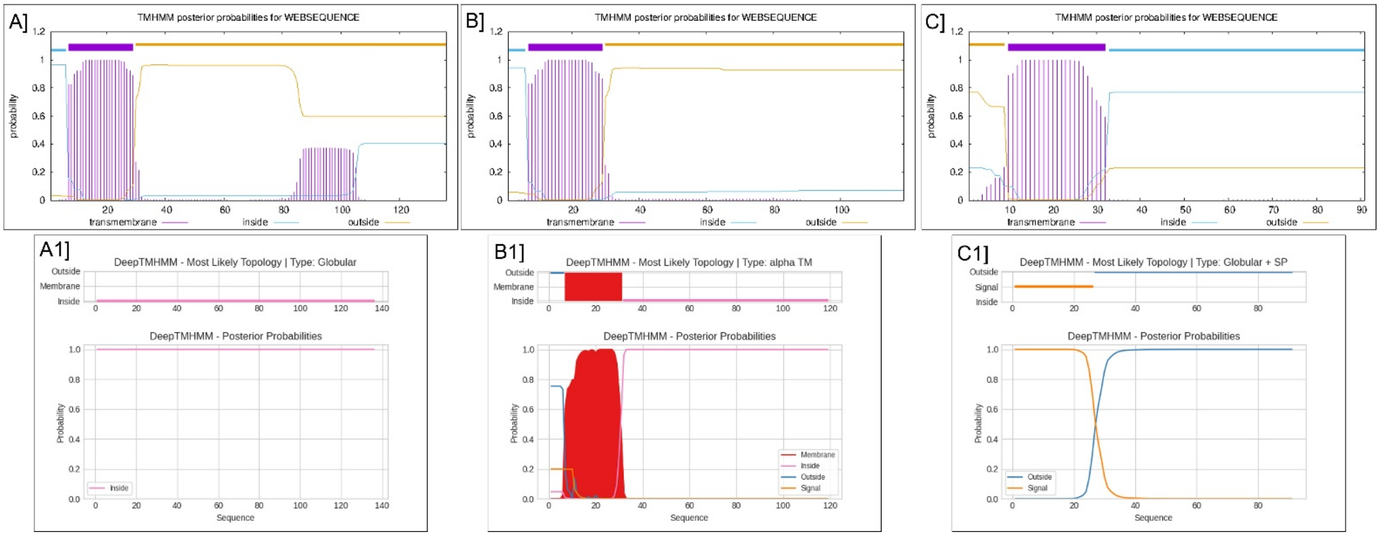


**Supplementary Figure 2: Problematic TMD topology prediction for Sgl$^{\phi \mathrm{Cb5}}$.** The graphical output of TMHMM and DeepTMHMM, showing Sgl$^{\phi \mathrm{Cb5}}$ and ϕX174 E-protein. (**A**) The lysis protein of ϕCb5 (136 residues) showing the probability of a TMD with N-in, C-out topology in TMHMM, and (**A1**) DeepTMHMM showing no TMD for Sgl$^{\phi \mathrm{Cb5}}$. (**B**) The truncated lysis protein of ϕCb5 (119 residues long, i.e., residues 89-105 removed) showing the probability of a TMD with N-in, C-out topology in TMHMM, and (**B1**) DeepTMHMM showing the probability of a TMD with N-out, C-in topology for truncated Sgl$^{\phi \mathrm{Cb5}}$. (**C**) The E-protein of ϕX174 showing N-out, C-in topology with one TMD in TMHMM, and (**C1**) DeepTMHMM predicts absence of TMD in favor of an N-terminal signal sequence.

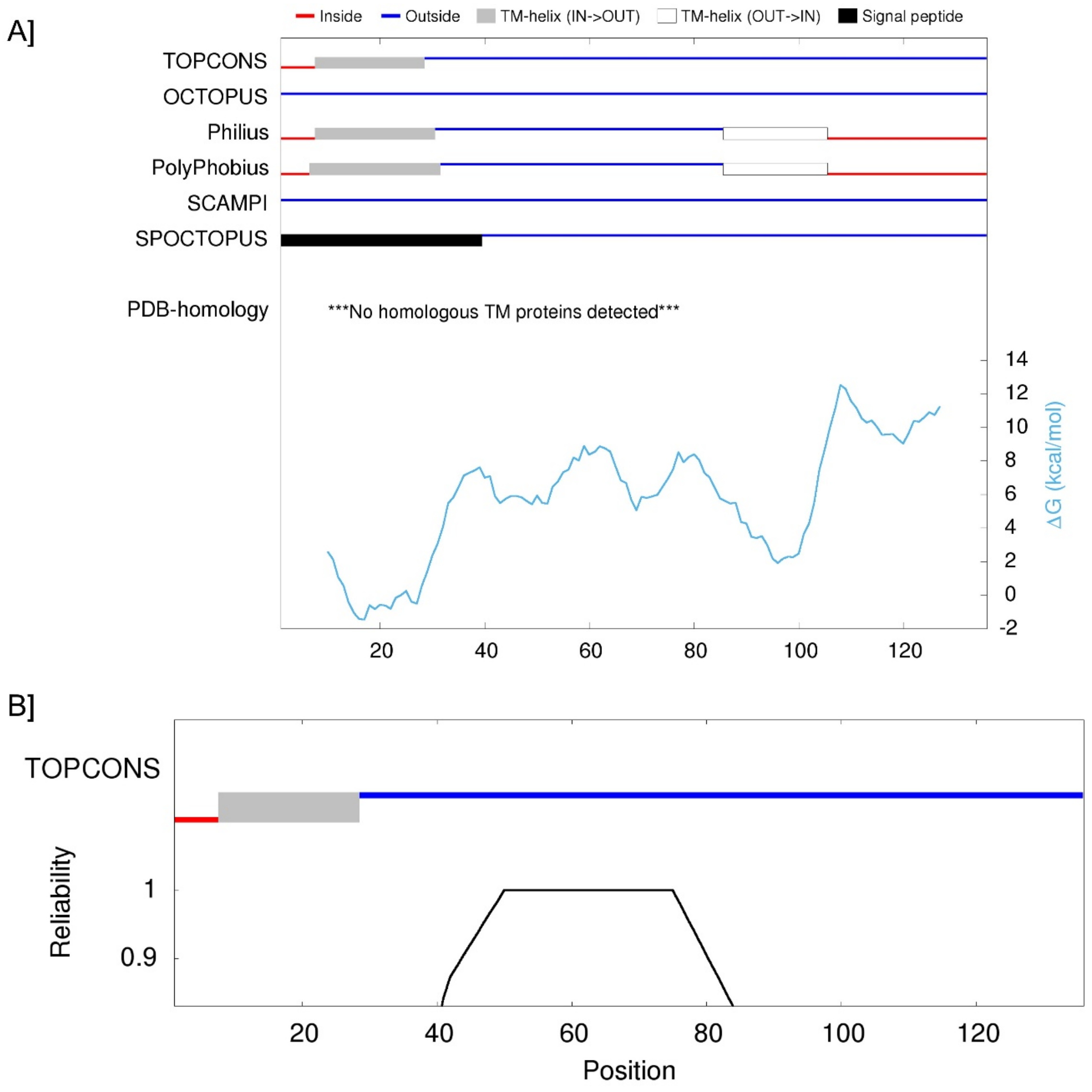


**Supplementary Figure 3: The graphical output of TOPCONS showing Sgl$^{\phi Cb5}$ topology.** (**A**) The output includes results from a variety of TMD prediction tools, and (**B**) overall consensus indicates that Sgl$^{\phi Cb5}$ has an N-in/C-out topology with one predicted TMD.

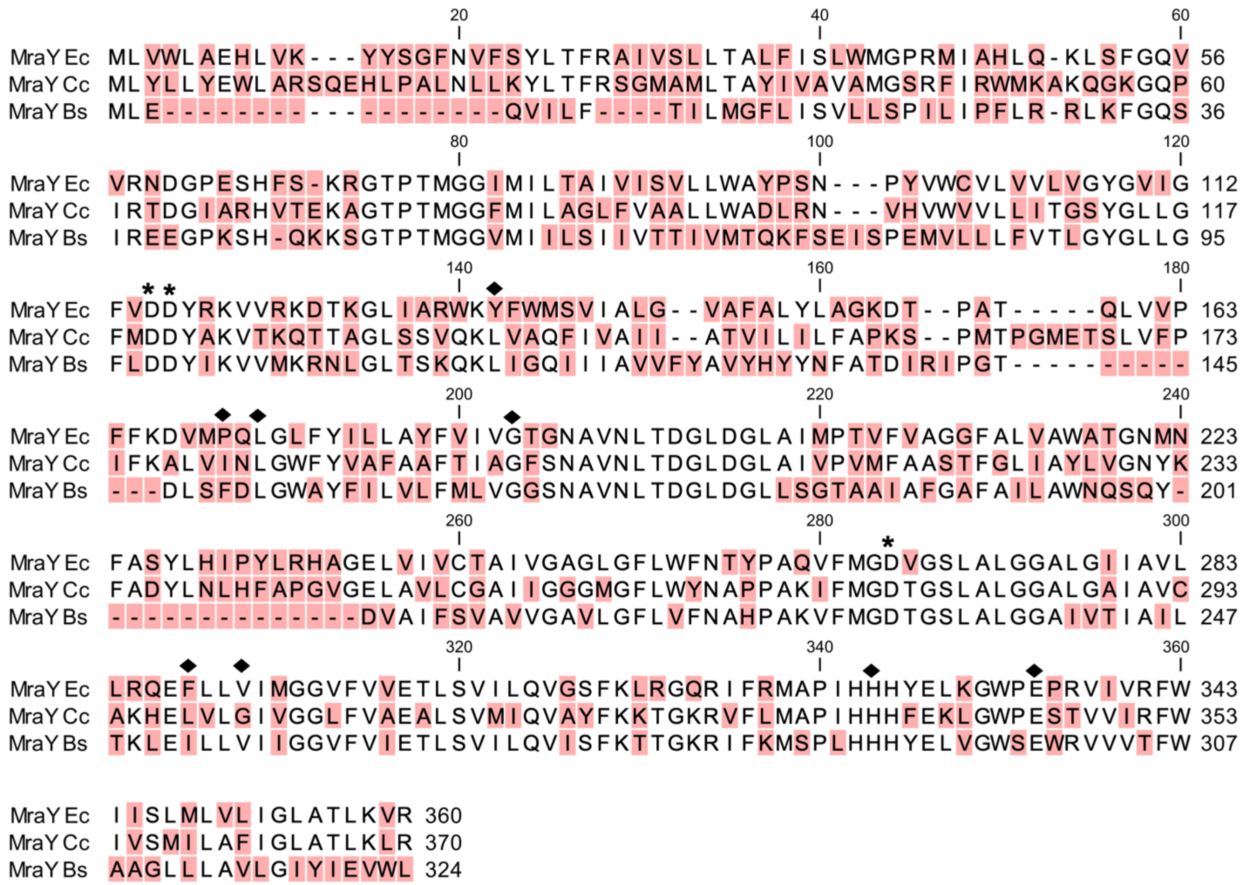


**Supplemental Figure 4: Alignment of MraY from *E. coli*, *C. crescentus* and *B. subtilis*.** The active site Asp residue is marked with an asterisk, and the diamond (black) represents the amino acids where E resistance mutations were identified previously (Orta et al., 2023). % identity between *E. coli* MraY and *C. crescentus* MraY is 39%, between *E. coli* MraY and *B. subtilis* MraY is 35%, and between *C. crescentus* MraY and *B. subtilis* MraY is 33%.

**A] *C. vibrioides* FKBP-type PPIase CC2799 [AAK24763.1]**

MVSLPSGLQYKVIASGPKDGPSPKLGDIIKVNYEGKLLSGAVFDSSFARGKAAIMPADGL
IRGWLEALPLMKVGDEWTLFIPPELGYGSRDMGEIPPDSTLVFRLKLEGMLSVD

**B] *C. vibrioides* FKBP-type PPIase CC3636 [AAK25598.1]**

MLAGRRILIEVPMHRRALLVTLAAAGVALSACGPSKKAQENLAAADAFMAKNAKEPGVVTLPQGLQYKVV
REGPNGGMHPNKADEVKVHYEGKLIDGTVFDSSYERGVPAVFPLDGLVPAWVIALQRMKAGDEWILYVPP
ALGYGAQDKGPIPGNSVMIFRIELLDVNRIGPGKPKE

**Supplementary Figure 5: Amino acid sequences of *C. vibrioides* FKBP-type PPIase; (A) CC2799 and (B) CC3636.** CC2799 is cytoplasmic, and CC3636 is periplasmic with an N-terminal signal sequence (highlighted). The genes were cloned into pZA32.

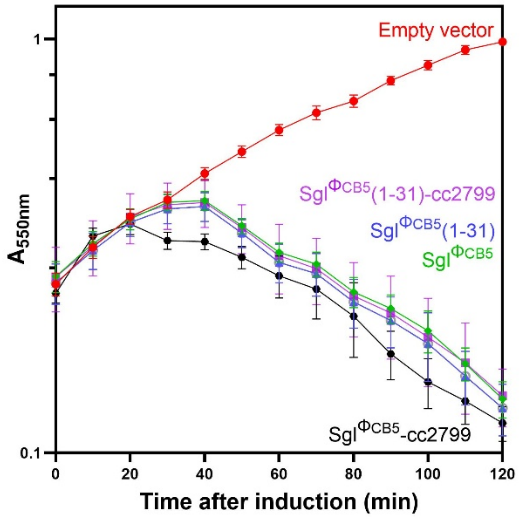


**Supplementary Figure 6: Effect of CC2799 expression on lysis mediated by full-length and truncated Sgl$^{\phi Cb5}$.** Sgl$^{\phi Cb5}$-mediated lysis in TB28 Ω *mraY$_{Cc}$*. Sgl$^{\phi Cb5}$ was cloned into pBAD24 and expressed in the TB28 Ω *mraY$_{Cc}$*-kan background at 37°C. Cultures were grown at an $A_{550}$ of ~0.2 and induced with arabinose at a final concentration of 0.4% (wt/vol). Lysis profile of truncated Sgl$^{\phi Cb5\ (1\text{-}31)}$ in the TB28 Ω *mraY$_{Cc}$*-kan co-transformed with pBAD24 *sgl$^{\phi Cb5}$* and compatible plasmid (pZA32) expressing FKBP-like PPIase CC2799. Empty vector (red), pBAD24 *sgl$^{\phi Cb5}$* + pZA32 empty (green), pBAD24 *sgl$^{\phi Cb5\ (1\text{-}31)}$* + pZA32 empty (blue), pBAD24 *sgl$^{\phi Cb5}$* + pZA32 CC2799 (black) and pBAD24 *sgl$^{\phi Cb5\ (1\text{-}31)}$* + pZA32 CC2799 (purple). The data represent the mean ± standard deviation of three independent experiments.

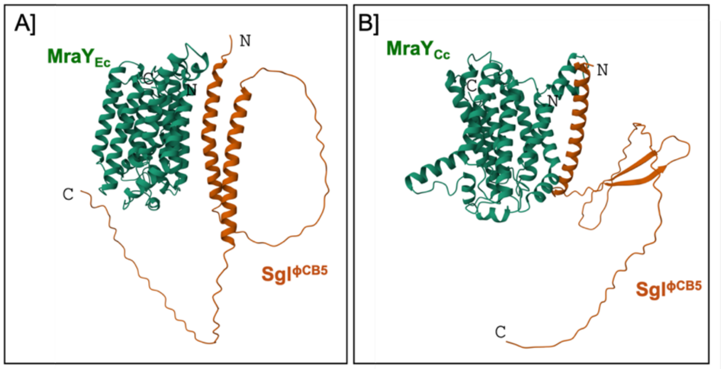


| **A. Model** | **MraY$_{Ec}$- Sgl$^{\phi Cb5}$ ipTM** | **Minimum inter-chain PAE (Å)** | **Overall pTM** |
|---|---|---|---|
| 0 | 0.19 | 15.39 | 0.68 |
| 1 | 0.17 | 15.78 | 0.68 |
| 2 | 0.14 | 15.77 | 0.67 |
| 3 | 0.13 | 15.46 | 0.68 |
| 4 | 0.12 | 16.44 | 0.68 |
| **B. Model** | **MraY$_{Cc}$- Sgl$^{\phi Cb5}$ ipTM** | **Minimum inter-chain PAE (Å)** | **Overall pTM** |
| 0 | 0.13 | 19.57 | 0.68 |
| 1 | 0.11 | 19.93 | 0.68 |
| 2 | 0.11 | 20.89 | 0.68 |
| 3 | 0.11 | 20.56 | 0.68 |
| 4 | 0.11 | 20.79 | 0.68 |

**Supplementary Figure 7: AlphaFold 3 predictions of MraY-Sgl$^{\phi Cb5}$ complexes.**
(**A**) MraY$_{Ec}$-Sgl$^{\phi Cb5}$ predictions; (**B**) MraY$_{Cc}$-Sgl$^{\phi Cb5}$ predictions, with the corresponding confidence metrics.
The table summarizes the predicted interface template modelling (ipTM) scores, minimum interchain predicted aligned error (PAE), and overall predicted template model ing (pTM) scores for five models each of MraY$_{Ec}$-Sgl$^{\phi Cb5}$ and MraY$_{Cc}$-Sgl$^{\phi Cb5}$.
The models are shown only as exploratory structural predictions and were not used to assign residue-level interactions.

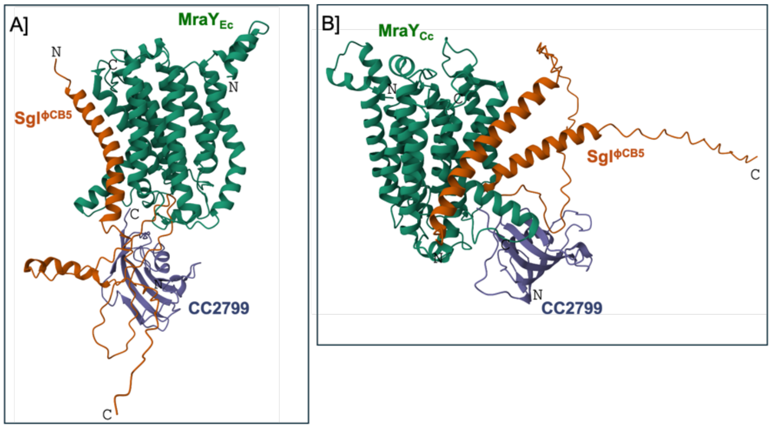


| Model | MraY$_{Ec}$- Sgl$^{\phi Cb5}$ - CC2799 ipTM | Overall pTM |
|---|---|---|
| 0 | 0.17 | 0.53 |
| 1 | 0.15 | 0.52 |
| 2 | 0.14 | 0.51 |
| 3 | 0.16 | 0.52 |
| 4 | 0.12 | 0.52 |
| **Model** | **MraY$_{Cc}$- Sgl$^{\phi Cb5}$ - CC2799 ipTM** | **Overall pTM** |
| 0 | 0.37 | 0.62 |
| 1 | 0.38 | 0.62 |
| 2 | 0.37 | 0.62 |
| 3 | 0.34 | 0.61 |
| 4 | 0.34 | 0.60 |

**Supplementary Figure 8: AlphaFold 3 predictions of MraY-Sgl$^{\phi Cb5}$ -CC2799 ternary complexes.** (**A**) MraY$_{Ec}$-Sgl$^{\phi Cb5}$-CC2799 predictions; (**B**) MraY$_{Cc}$-Sgl$^{\phi Cb5}$-CC2799 predictions, with the corresponding confidence metrics.
The table summarizes the overall ipTM and pTM scores for five predicted models each of MraY$_{Ec}$-Sgl$^{\phi Cb5}$-CC2799 and MraY$_{Cc}$-Sgl$^{\phi Cb5}$-CC2799.
The ternary predictions do not confidently establish a stable three-protein complex. Therefore, models are shown only as exploratory structural predictions rather than evidence that CC2799 bridges Sgl$^{\phi Cb5}$ and MraY.

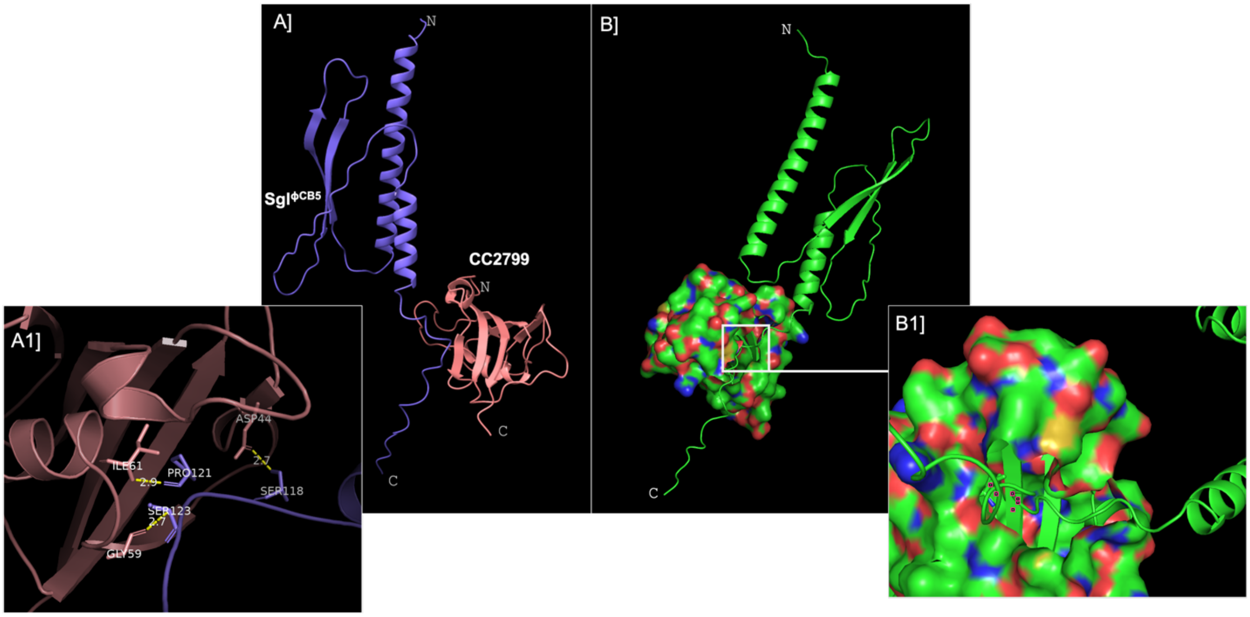


**Supplementary Figure 9: AlphaFold 3-predicted structural model of Sgl$^{\phi Cb5}$ in association with the FKBP-like CC2799. (A)** Cartoon representation of the predicted Sgl$^{\phi Cb5}$–CC2799 complex visualized using ChimeraX. **(A1)** Enlarged view of the predicted interface highlighted in panel A, showing selected residues from CC2799 (Asp44, Gly59, and Ile61) and Sgl$^{\phi Cb5}$ (Ser118, Pro121, and Ser123). Yellow dashed lines indicate interatomic distances between the displayed residues, with values shown in Å. **(B)** Alternative visualization of the predicted Sgl$^{\phi Cb5}$–CC2799 complex generated using PyMOL. Sgl$^{\phi Cb5}$ is shown as a green cartoon, while CC2799 is represented as a molecular surface colored by atom type. The model illustrates the predicted proximity of the Sgl$^{\phi Cb5}$ C-terminal region to the CC2799 surface. **(B1)** Magnified view of the putative interface indicated by the white box in panel B, highlighting the placement of the Sgl$^{\phi Cb5}$ polypeptide near the CC2799 molecular surface. The predicted protein–protein interface has low confidence; these models should be interpreted as exploratory and do not establish direct binding.

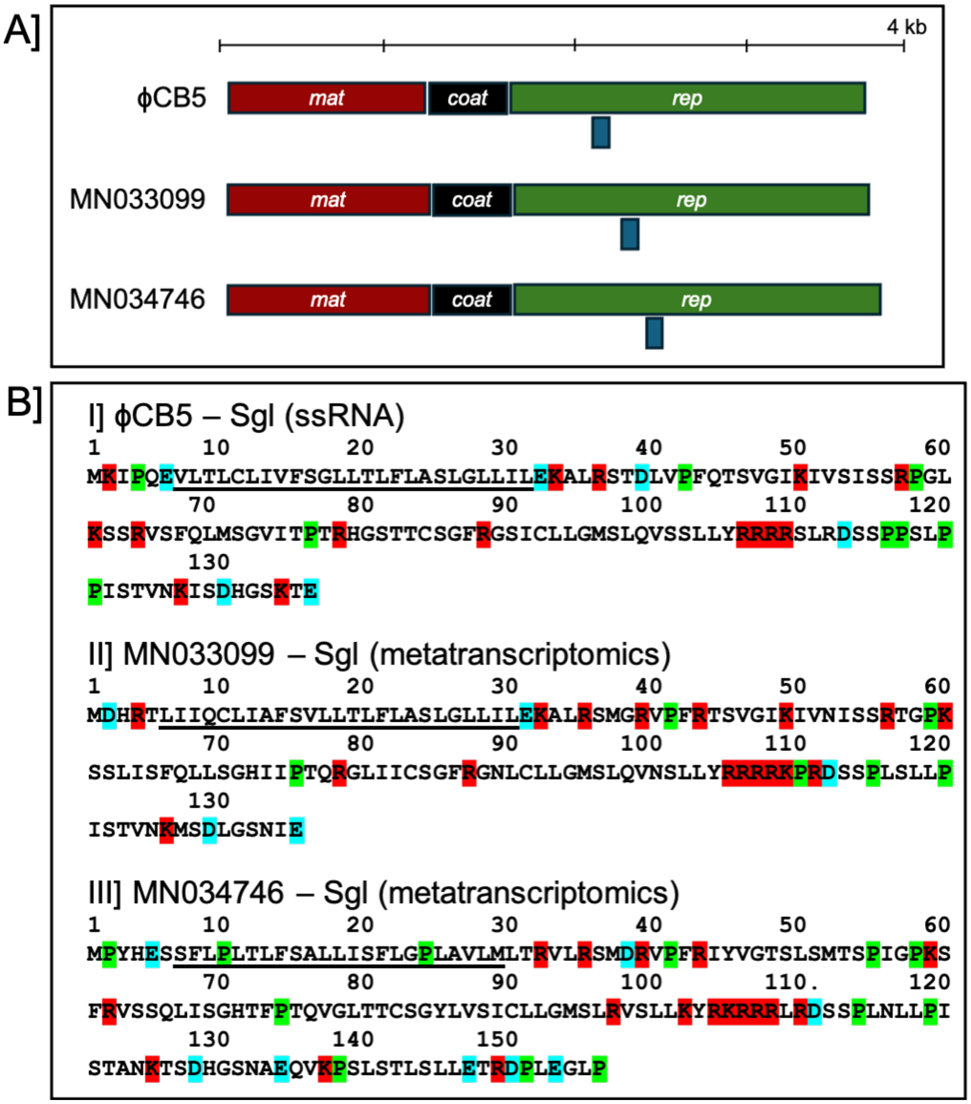


**Supplementary Figure 10: Comparative genome organization and distribution of charged residues in putative Sgl proteins of ϕCb5-related ssRNA phages.** (**A**) Schematic representation of the genomic organization of three ϕCb5-related ssRNA phages, showing the maturation (*mat*), coat (*coat*), and replicase (*rep*) genes. The short blue boxes beneath the *rep* genes indicate the positions of the predicted Sgl open reading frames. (**B**) Comparison of the distribution of selected amino acid residues within the three predicted Sgl proteins. Acidic and basic residues are blue and red, respectively. Proline (P) residues are shown in green. Underline denotes TMD.

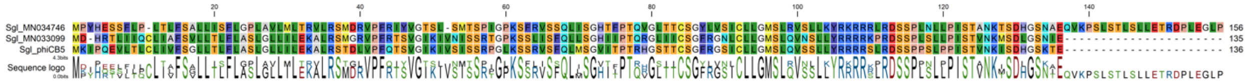


**Supplementary Figure 11: Multiple sequence alignment of Sgl$^{\phi Cb5}$ and two putative homologs.** We aligned the amino acid sequences of Sgl$^{\phi Cb5}$ and two putative homologs identified from GenBank accessions MN033099 and MN034746 to assess sequence conservation. The alignment highlights identical and chemically similar residues using the alignment colour scheme, and the conservation histogram indicates the degree of sequence similarity across aligned positions. Conserved residues and sequence motifs are distributed throughout the proteins, including the C-terminal region. The alignment supports sequence-level relatedness but does not establish that the putative homologs possess lytic activity or share the same molecular target.

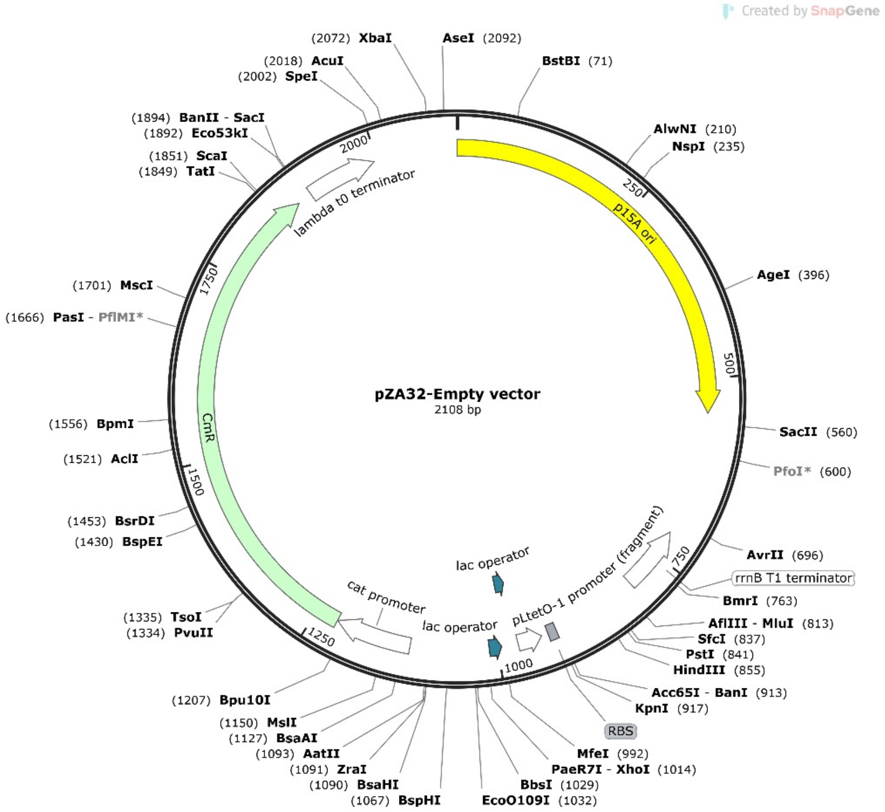


**Supplementary Figure 12: Genetic map of the pZA32 empty vector (2,108 bp).** The plasmid contains the p15A origin of replication, chloramphenicol resistance gene (CmR), PLtetO-1 promoter, tandem lac operator sequences, ribosome-binding site (RBS), rrnB T1 transcription terminator, and λ t0 terminator. Restriction enzyme recognition sites are indicated around the plasmid circumference. The vector was used as the empty-vector control in all expression experiments.
